# Plant-specific histone deacetylases are essential for early as well as late stages of Medicago nodule development

**DOI:** 10.1101/2020.11.09.374819

**Authors:** Huchen Li, Stefan Schilderink, Qingqin Cao, Olga Kulikova, Ton Bisseling

**Affiliations:** Beijing Advanced Innovation Center for Tree Breeding by Molecular Design, Beijing University of Agriculture, Beijing 102206, China; Department of Plant Sciences, Laboratory of Molecular Biology, Wageningen University, Droevendaalsesteeg 1, 6708 PB Wageningen, The Netherlands; College of Plant Science and Technology, Beijing Key Laboratory for Agricultural Application and New Technique, Beijing University of Agriculture, Beijing 102206, China; St. Bonifatius college, Burgemeester Fockema Andreaelaan 7-9, 3582 KA Utrecht, The Netherlands

## Abstract

Legume and rhizobium can establish a nitrogen-fixing nodule symbiosis. Previous studies have shown that several transcription factors that play a role in (lateral) root development are also involved in nodule development. Chromatin remodelling factors, like transcription factors, are key players in regulating gene expression. However, it has not been studied whether chromatin remodelling genes that are essential for root development get involved in nodule development. Here we studied the role of Medicago histone deacetylases (MtHDTs) in nodule development. Their Arabidopsis orthologs have been shown to play a role in root development. The expression of *MtHDTs* is induced in nodule primordia and is maintained in nodule meristem and infection zone. Conditional knock-down of their expression in a nodule-specific way by RNAi blocks nodule primordium development. A few nodules still can be formed but their nodule meristems are smaller and rhizobial colonization of the cells derived from the meristem is markedly reduced. Although the HDTs are expressed during nodule and root development, transcriptome analyses indicate that HDTs control the development of these organs in a different manner. During nodule development the MtHDTs positively regulate *3-hydroxy-3-methylglutaryl coenzyme a reductase 1* (*MtHMGR1*). The decreased expression of *MtHMGR1* is sufficient to explain the block of primordium formation.

**ONE SENTENCE SUMMARY:** Plant-specific histone deacetylases regulate the expression of *3-hydroxy-3-methylglutaryl-coenzyme A reductases* to control root nodule development.

## INTRODUCTION

Plants are able to develop lateral organs post-embryonically. An example is the formation of lateral roots (Malamy and Benfey, 1997). Roots of legume plants have the property to form a second lateral organ, root nodules. The latter are symbiotic organs which are used to host rhizobium bacteria. These become able to reduce atmospheric nitrogen into ammonia which can be used by the host (Udvardi and Poole, 2013).

The model legume Medicago (*Medicago truncatula*) forms indeterminate nodules. Their histology and ontology bear resemblance to that of (lateral) roots. In both organs a meristem is present at their apex (Franssen et al., 1992; van den Berg et al., 1995), which is followed by a zone containing differentiating cells. This is the elongation zone in roots and the infection zone in nodules (Vinardell et al., 2003; Vanstraelen et al., 2009). In the latter intracellular infection by rhizobia takes place. The fully differentiated cells form the differentiated zone in roots and the fixation zone in nodules. The switch from infection to fixation zone is characterized by the sudden accumulation of starch in the infected cells (Gavrin et al., 2014). In Medicago, both nodules and lateral roots are developed from primordia whose formation is initiated at the protoxylem pole and starts with cell division in pericycle and subsequently divisions are induced in endodermis and cortex in both cases (Dubrovsky et al., 2001; Xiao et al., 2014; Xiao et al., 2019). Therefore nodules and lateral roots show similarities in organogenesis.

Recent studies showed that some transcription factors involved in (lateral) root development have been recruited for nodule development. In Medicago, knock-down of *PLETHORA* genes known to be key regulators in root development, blocks nodule meristem activity (Aida et al., 2004; Franssen et al., 2015), and knock-out of *LOB-DOMAIN PROTEIN 16* (*LBD16*) reduces both nodule and lateral root initiation (Goh et al., 2012; Schiessl et al., 2019). It is known that chromatin remodelling factors contribute to transcriptional reprogramming and also play a central role in plant organ development (Jarillo et al., 2009). However, whether chromatin remodelling factors which are involved in root development, also have a role in nodule development has never been studied.

Previously, we have shown that in Arabidopsis two plant-specific histone deacetylases (*AtHDT1/2*) are expressed in the root meristem, and control its size by repressing *C*_*19*_*-GIBBERELLIN 2-OXIDASE 2* (*AtGA2ox2*) (Li et al., 2017). Further, *AtHDTs* are markedly up-regulated in dedifferentiating pericycle cells during the initiation of lateral root primordia (De Smet et al., 2008). Medicago contains 3 *HDT* members; *Medtr4g055440*, *Medtr2g084815* and *Medtr8g069135*, they were designated as *MtHDT1*, *MtHDT2 and MtHDT3*, respectively (Grandperret et al., 2014). Laser capture microdissection RNA sequencing (LCM-RNA-seq) analyses indicated that they all are expressed in nodule meristem and infection zone (Roux et al., 2014). Here we studied whether Medicago HDTs play a role in nodule development, and if so whether they have a similar function as in the root development.

We showed that the 3 *MtHDTs* are expressed in young nodule primordia. In mature nodules they are expressed in the meristem and the infection zone. Knock-down of *MtHDTs* in a nodule specific way (*ENOD12::MtHDTs RNAi*) blocks cell division in most of the nodule primordia. In the few nodules formed on RNAi transgenic roots, meristem size and activity, as well as rhizobial colonization are reduced. Transcriptome analysis of RNAi nodules showed that HDTs regulate nodule and root development in a different manner. The differentially expressed genes in RNAi nodule primordia and in mature nodules are in part overlapped, and in both cases expression of the *MtHMGR1* is reduced.

## RESULTS

### Medicago HDT2 Has A Similar Function as Arabidopsis HDT1/2 in Controlling Root Development

To compare the functions of the Medicago HDTs with the previously characterized Arabidopsis HDTs, we first analysed the phylogenetic relationship of HDTs by using protein sequences from several dicots and the monocot rice. This showed that HDTs in rice were separated from those in dicots. Within dicots HDTs have evolved into two clades (Fig. 1A, Supplemental Table S1). The first clade contained the Arabidopsis AtHDT3 and none of the Medicago MtHDTs. The second clade contained AtHDT1, 2, 4 and all 3 MtHDTs. Further, independent duplications have occurred in the 3 legume species, Medicago, Lotus and Soybean, and this have resulted in highly homologous HDT pairs. In Medicago such pair is formed by MtHDT2 and 3. In Arabidopsis a similar independent duplication resulted in AtHDT1 and 2. Previously, we showed that AtHDT1 and 2 are functionally redundant and are essential for root growth. AtHDT4 regulates root growth as well (Han et al., 2016). Therefore it is very likely that also some of the MtHDTs are involved in root development.

**Figure 1.**
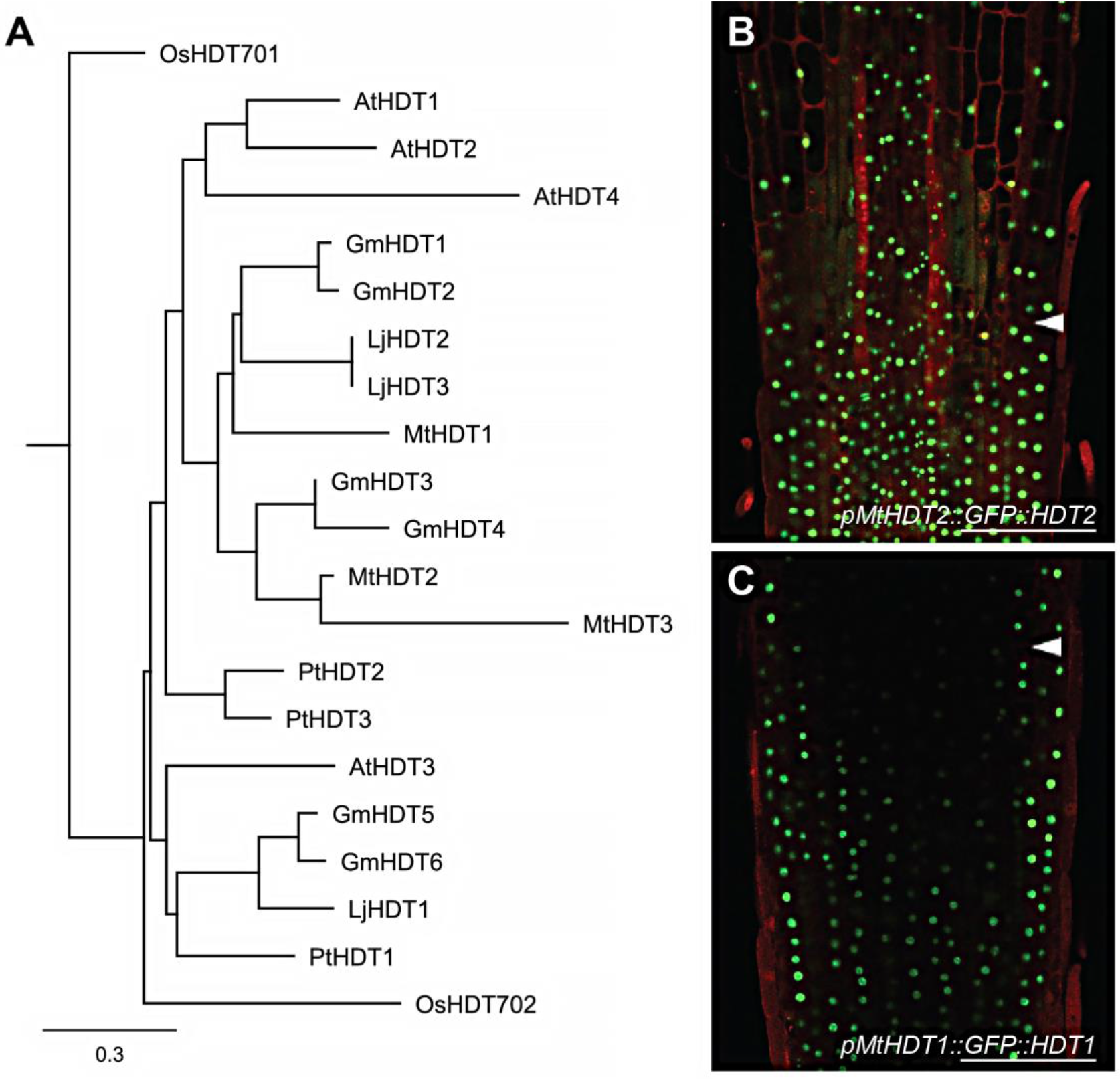
MtHDTs are orthologous to AtHDT1, 2. A, Phylogenetic tree of HDT proteins. The protein sequences are obtained from *Medicago truncatula* (Mt), *Lotus japonicus* (Lj), *Glycine max* (Gm), *Arabidopsis thaliana* (At), *Populus trichocarpa* (Pt) and *Oryza sativa* (Os). Scale bar represents substitution per site. B and C, Localization of *pMtHDT2::GFP::HDT2* (B) and *pMtHDT1::GFP::HDT1* (C) in longitudinal sections of Medicago root tips. Arrowheads indicate the boundary between root meristem and elongation zone. GFP signal is localized in nuclei. Scale bar=100μm.

To study whether MtHDTs have a similar expression pattern as AtHDTs in roots, we generated *GFP-MtHDT* constructs including a ~2kb DNA region upstream of the start codon (putative promoter), GFP and the corresponding *MtHDT* coding sequence (*pMtHDT1::GFP::HDT1*, *pMtHDT2::GFP::HDT2* and *pMtHDT3::GFP::HDT3*). These constructs were introduced into Medicago by *Agrobacterium rhizogenes* mediated hairy root transformation (Limpens et al., 2004). In Medicago roots, *pMtHDT1::GFP::HDT1*and *pMtHDT2::GFP::HDT2* were expressed in the meristem and elongation zone and GFP fluorescence was mainly detected in nucleoli (Figs. 1, B and C). In the differentiated zone these fusion proteins were hardly detected. This is similar to the expression pattern and the subcellular localization of AtHDT1 and AtHDT2 in Arabidopsis root tips (Li et al., 2017). Expression level of MtHDT2 in root tips was higher than that of MtHDT1. Expression of *pMtHDT3::GFP::HDT3* was below detection level, therefore we studied the *MtHDT3* expression pattern using a *pMtHDT3::GUS* construct including the putative *MtHDT3* promoter and *β-glucuronidase (GUS)* coding sequence. The construct was introduced into Medicago by hairy root transformation and it showed that *pMtHDT3::GUS* was weakly expressed in the root meristem (Supplemental Fig. S1).

The high homology and the similar expression pattern of HDTs in Arabidopsis and Medicago roots suggests that they may control root growth in the same way. A *Mthdt2* Tnt1 mutant containing mutations either in the third exon or in the eighth intron has recently become available, but it has a wild-type like root phenotype (Supplemental Fig. S2) and for the other HDT genes mutants are not available. To determine which *MtHDT* gene is sufficient to support root growth in Arabidopsis, we introduced each *pMtHDT::GFP::HDT* construct into a double heterozygous *HDT1hdt1HDT2hdt2* Arabidopsis mutant. Loss of function of both *AtHDT1* and *AtHDT2* is lethal (Li et al., 2017), therefore we tested in the progeny of the transformed *HDT1hdt1HDT2hdt2* plants which *MtHDT* gene was able to rescue the lethal phenotype. More than 200 transformed plantlets of each progeny were genotyped, this showed that *pMtHDT2::GFP::HDT2* complemented Arabidopsis *hdt1hdt1hdt2hdt2*, whereas *pMtHDT1::GFP::HDT1* and *pMtHDT3::GFP::HDT3* did not. Further, in Arabidopsis roots *pMtHDT2::GFP::HDT2* was expressed in the meristem and elongation zone and mainly localized in nucleoli (Supplemental Fig. S3), similar to AtHDT1/2 (Li et al., 2017). The expression pattern studies and complementation test together suggest that MtHDT2 has a similar role as AtHDT1, 2 in root development. It does not exclude that MtHDT1 and 3 are also involved in root development as they are expressed in Medicago root tips.

### MtHDTs Are Expressed in the Nodule Meristem and Infection Zone

In this study we especially focused on the role of MtHDTs in nodule development. As all 3 Medicago HDTs are expressed in roots, and nodule and root development are related, we studied first all 3 Medicago genes. To determine where *MtHDTs* are expressed in nodules, we performed RNA *in situ* hybridisation on longitudinal sections of nodules using probe sets specific for each *MtHDT*. We used *in situ* hybridisation as this gives the most accurate expression pattern, especially since we could not test in Medicago whether the selected *MtHDT* promoter regions are biologically functional. The *in situ* hybridisation experiment showed that *MtHDT2* transcripts were present at a similar level in both the meristem and infection zone (Fig. 2A). In the latter, *MtHDT2* was mainly expressed in infected cells and hardly detectable in uninfected cells. This is different from roots in which *HDT* genes are only expressed in the meristem (Li et al., 2017). At the transition from infection to fixation zone, the expression of *MtHDT2* dropped dramatically. The spatial distribution of *MtHDT1* and *MtHDT3* transcripts was similar to that of *MtHDT2*, but the hybridisation signals were markedly lower (Figs 2, C and D). So like in roots, *MtHDT2* is higher expressed in nodules than the other *MtHDTs.* In addition, *MtHDT2* is certainly involved in root development. Therefore in further experiments we focused on this gene.

**Figure 2.**
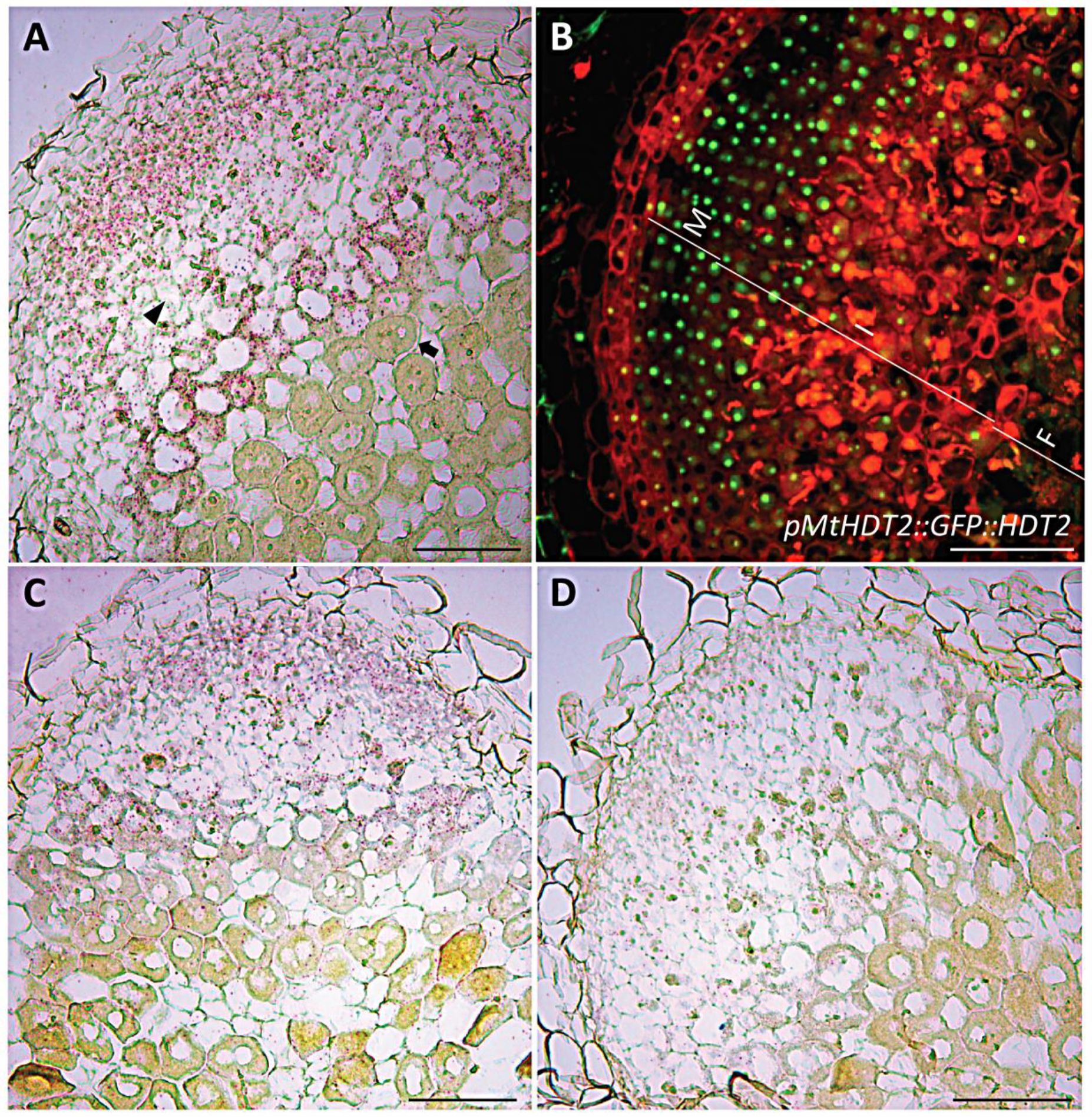
*MtHDTs* are expressed in the nodule meristem and infection zone. A, Expression of *MtHDT2* mRNA visualized by *in situ* hybridisation in wild-type Medicago nodules. The arrowhead indicates a non-infected cell in the infection zone, the arrow indicates a cell of the first cell layer of the fixation zone where amyloplasts are detectable at the periphery. B, Localization pattern of *pMtHDT2::GFP::HDT2* in nodules. The nodule meristem zone (M), infection zone (I) and fixation zone (F) are marked. C and D, Expression of *MtHDT1* (C) and *MtHDT3* (D) mRNA visualized by *in situ* hybridisation in wild-type Medicago nodules. Images are longitudinal sections of nodules harvested at 21dpi. Representative image is shown. In A, C and D, red dots are hybridisation signals. Scale bar=100μm.

To determine the subcellular localization of MtHDT2 protein in nodules, we used *pMtHDT2::GFP::HDT2* construct. This showed that MtHDT2 protein accumulated in cells of nodule meristem and infection zone, and like in roots, mainly in nucleoli. Further, at the switch from infection to fixation zone its level suddenly dropped to below detection level (Fig. 2B). So the distribution of the protein is similar to that of the transcript. Further, the expression of *MtHDT2* in meristem and infected cells of the infection zone indicated that this gene might control meristem activity, rhizobial release and/or intracellular accommodation of rhizobia.

### Meristem Activity and Probably Rhizobial Colonization Require MtHDTs

To determine the role of MtHDTs in nodules, we made a nodule-specific RNA interference construct to target all 3 *MtHDT* transcripts (*ENOD12::MtHDTs RNAi*). Although *MtHDT2* has the highest expression level in nodules we decided also to knock-down the other 2 *MtHDTs*, as a *MtHDT2* Tnt1 mutant has no nodule phenotype (Supplemental Fig. S2). We used the *ENOD12* promoter to drive the RNAi construct as it is active in the nodule meristem and infection zone and so it covers the expression domains of the 3 *MtHDTs* (Limpens et al., 2005; Franssen et al., 2015). In the RNAi transgenic nodules *MtHDT1, 2* and *3* were knocked-down to 22%, 7% and 29% of the levels in *ENOD12-EV* (**E**mpty **V**ector) control nodules, respectively (Fig. 3A). At 21 days post inoculation (dpi), control roots formed on average 6.0 nodules/root, whereas *MtHDTs RNAi* roots had only 1.1 nodules/root (Fig. 3B). Although the RNAi nodule number was low, it still allowed their histological characterization.

**Figure 3.**
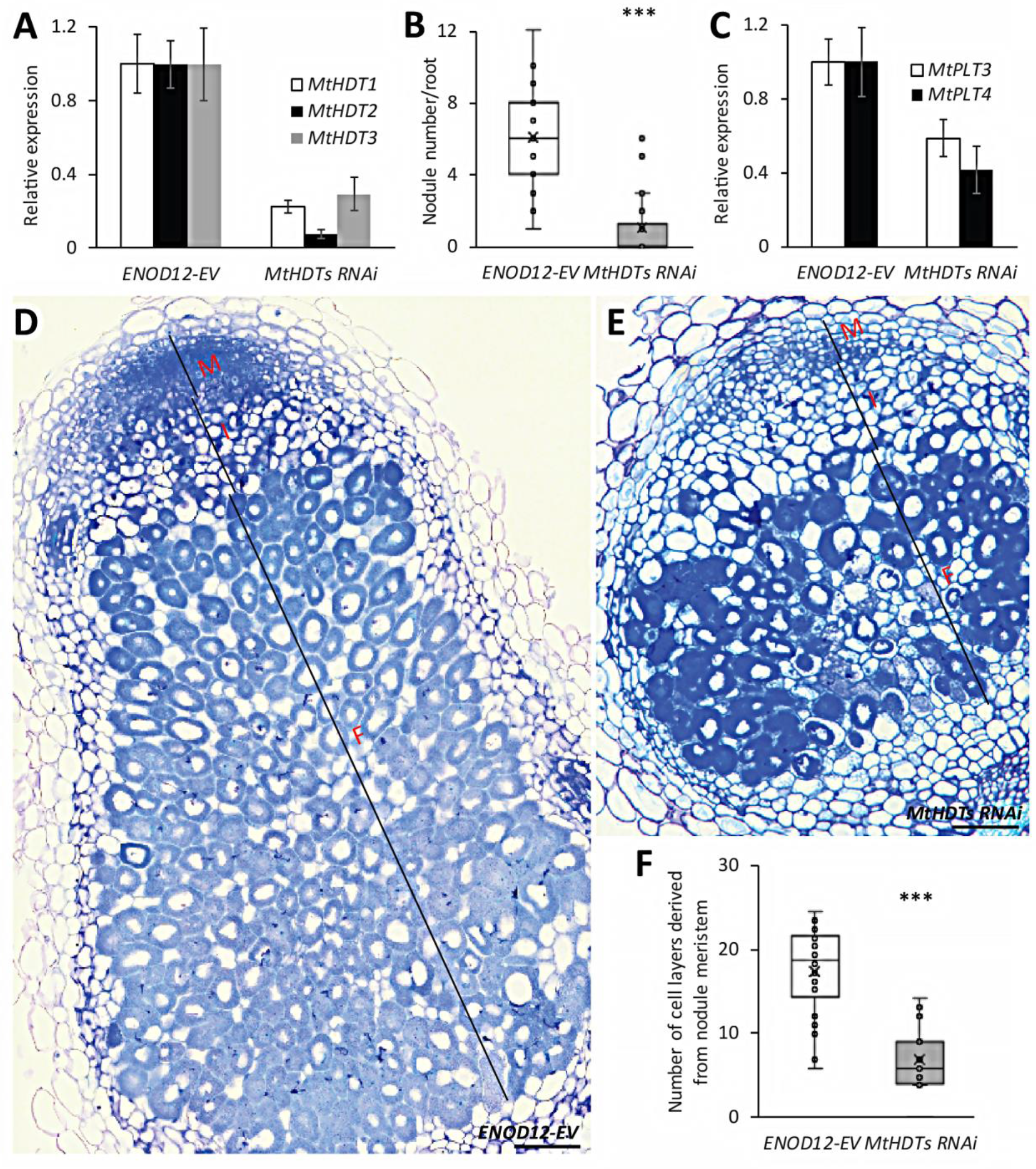
Knock-down of *MtHDTs* affects nodule meristem functioning and rhizobial colonization. A, Reverse transcription quantitative PCR (RT-qPCR) analysis of *MtHDTs* expression in *ENOD12-EV* control and *MtHDTs RNAi* nodules. B, Nodule number formed per *ENOD12-EV* and *MtHDTs RNAi* transgenic root (n>20). C, RT-qPCR analysis of *MtPLT3, 4* expression in *ENOD12-EV* control and *MtHDTs RNAi* nodules. D and E, Morphology of *ENOD12-EV* (D) and *MtHDTs RNAi* (E) nodules studied by light microscopy. Representative longitudinal sections are shown. The nodule meristem (M), infection zone (I) and fixation zone (F) are marked. Scale bar=100μm. F, Number of cell layers derived from nodule meristem in *ENOD12-EV* and *MtHDTs RNAi* transgenic nodules (n>15). Nodules were harvested at 21dpi. Panels in A and C show mean±SEM determined from three independent experiments. Asterisks in B and F indicate significant differences (***, p<0.001; Student’s *t* test).

The control nodules were elongated, whereas *MtHDTs RNAi* nodules were spherical and markedly smaller (Figs 3, D and E). Longitudinal sections of control nodules (n=22) showed that meristems were present at the apex of all nodules and contained ~8 cell layers (Fig. 3D). Meristems were also present in *MtHDTs RNAi* nodules (n=20), but only had ~4 cell layers (Fig. 3E). In agreement with this reduced number of layers, expression of *MtPLT3* and *MtPLT4*, two genes that are expressed throughout the nodule meristem (Franssen et al., 2015), was reduced to 59% and 42% of the control level in *MtHDTs RNAi* nodules (Fig. 3C).

About 8 cell layers of the proximal part of the central tissue of a mature nodule are formed and infected at the primordium stage, and are not derived from the nodule meristem (Xiao et al., 2014). *MtHDTs RNAi* nodules had about 8 cell layers at the proximal part with fully infected cells. They were completely packed with elongated symbiosomes (Figs 3, D and E). This is similar to control nodules. However, the number of cell layers derived from the nodule meristem was markedly reduced (Fig. 3F). Further, in the infected cells in these layers the colonization level was rather low, resulting in cells with large vacuoles and few bacteria. Collectively, these data showed that in the *MtHDTs RNAi* nodules knock-down of *MtHDTs* reduced nodule meristem size, and it affected the rhizobial colonization process in cells derived from the meristem, but not from primordium cells.

### Knock-down of *MtHDTs* Affects Nodule Primordium Development

As nodule number was markedly reduced on the RNAi roots we assumed that nodule primordium formation was affected. To test this, we transformed Medicago *ENOD11::GUS* plants (Boisson-Dernier et al., 2005) with the *MtHDTs RNAi* and *ENOD12-E*V construct, respectively, by hairy root transformation. The *ENOD11* promoter is active in the whole young nodule primordia, and it is only expressed in 1 or 2 cell layers adjacent to root vasculature in lateral root primordia (Supplemental Fig. S4). So it facilitates to distinguish nodule and lateral root primordia and to accurately count nodule primordium number.

Rhizobia were spot inoculated at the susceptible zone of 110 transgenic *ENOD12-E*V and 110 *MtHDTs RNAi* roots with a similar length. After 5 days, 99 control and 102 *MtHDTs RNAi* inoculated roots formed nodule primordia expressing *ENOD11*. The inoculated root segments with nodule primordia (~0.3cm) were embedded in plastic and sectioned to study till which stage nodule primordia had developed. In case of root segments containing more than one primordium, only the largest nodule primordium was counted. We successfully characterized 87 and 86 control and *MtHDTs RNAi* segments, respectively. This showed that in control roots, 90% (78 out of 87) of nodule primordia passed stage II and a relatively high number of them (54%, 47 out of 87) developed into or passed stage V (Fig. 4A). In contrast, on *MtHDTs RNAi* transgenic roots, the majority of nodule primordia (59%, 51 out of 86) were in stage I or stage II, only few *MtHDTs RNAi* nodule primordia (7%, 6 out of 86) had developed into or passed stage V (Fig. 4A). This suggested that the development of the majority of *MtHDTs RNAi* nodule primordia was blocked at an early stage, which is consistent with reduced nodule number at 21dpi (Fig. 3B).

**Figure 4.**
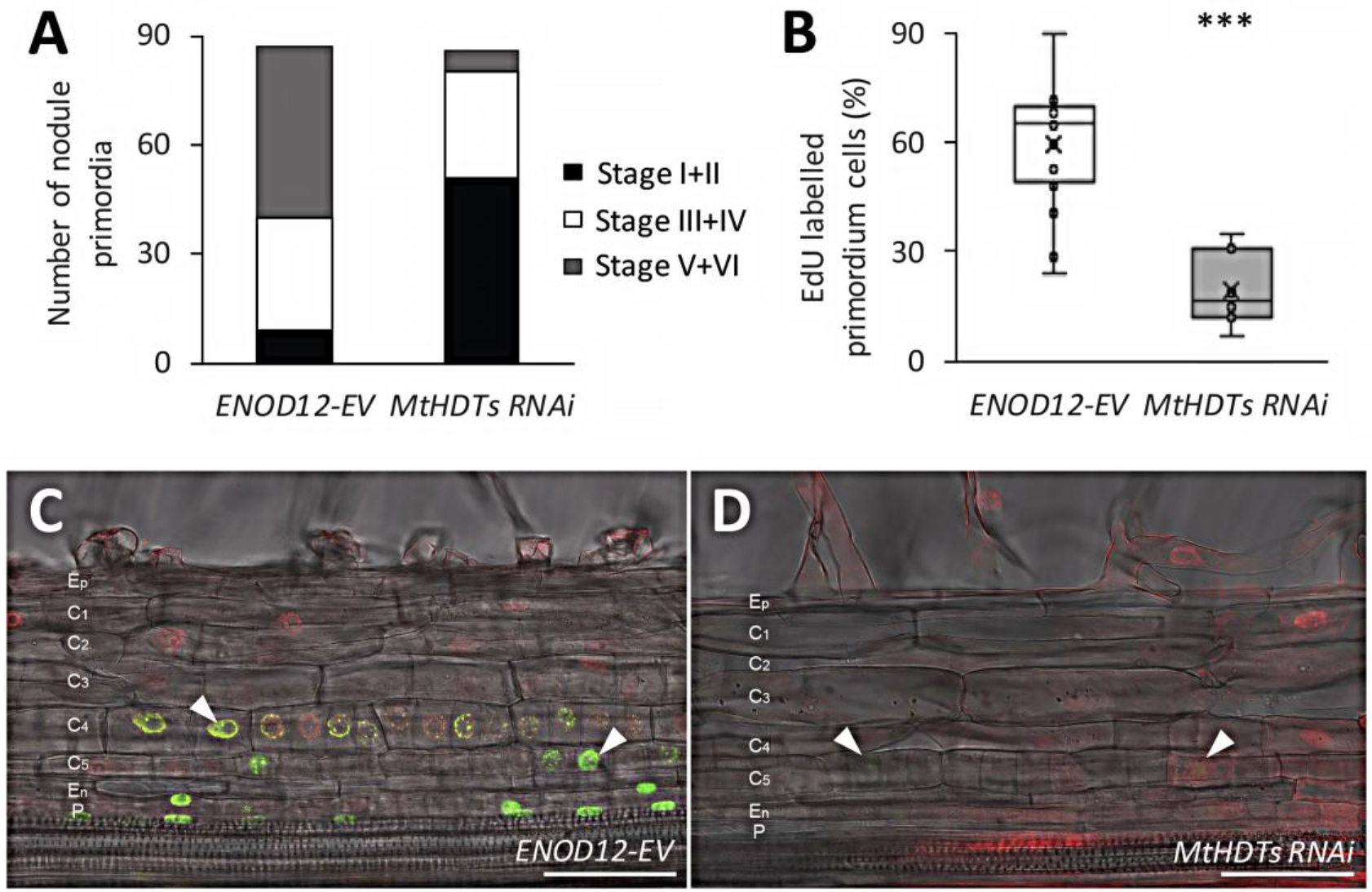
Knock-down of *MtHDTs* blocks nodule primordium development. A, Analysis of developmental stages of 5dpi *ENOD12-EV* (n=87) and *MtHDTs RNAi* (n=86) nodule primordia. B, Percentage of EdU labelled nodule primordium cells in 2dpi *ENOD12-EV* (n=15) and *MtHDTs RNAi* (n=7) nodule primordia. Nodule primordium cells were defined as divided or dividing cells that have smaller size. 8 *MtHDTs RNAi* nodule primordia have no EdU labelling and are not used for statistics. Asterisk indicates significant differences (***, p<0.001; Student’s *t* test). C and D, EdU signals in 2dpi *ENOD12-EV* (C) and *MtHDTs RNAi* (D) nodule primordia. Arrowheads indicate strong (C) or weak (D) green fluorescent signals in nuclei. Identical confocal microscope settings were used in C and D. P, Pericycle; En, Endodermis; C_5/4/3/2/1_, the fifth/fourth/third/second/first cortical cell layer; Ep, Epidermis. Scale bar=100μm.

To further support that *MtHDTs RNAi* nodule primordia were blocked in development, root segments containing nodule primordia were collected at 2 days after spot inoculation, they were then incubated for 2 hours with EdU, that is incorporated into replicating DNA during mitosis (Kotogany et al., 2010). By quantifying the percentage of EdU labelled nodule primordium cells, we could determine whether knock-down of *MtHDTs* reduced mitotic activity in young primordia. 15 control and 15 *MtHDTs RNAi* nodule primordia were analysed. All control nodule primordia had EdU labelled cells, and on average 62% of the primordium cells were labelled (Figs 4, B and C). In contrast, only 47% (7 out of 15) of *MtHDTs RNAi* nodule primordia had EdU labelled cells and in these primordia the percentage of labelled cells had markedly dropped to 20% (Figs. 4, B and D). Further, the intensity of fluorescence in EdU labelled cells was reduced in comparison with that in control primordia (Figs. 4, C and D). Therefore, we concluded that the development of the majority of *MtHDTs RNAi* nodule primordia had been blocked at the early stages.

### *MtHDTs* Are Expressed in Young Nodule Primordia

The block of *MtHDTs RNAi* nodule primordium development prompted us to study whether *MtHDTs* were expressed in nodule primordia. We first performed RNA *in situ* hybridisation for *MtHDT2*, as it has the highest expression level, on longitudinal sections of nodule primordia. Cell divisions in Medicago nodule primordia occur first in the pericycle and subsequently in the fifth cortical layer (C_5_) (Xiao et al., 2014). Such an early stage nodule primordium (stage I/II) is shown in Fig. 5A, *MtHDT2* transcripts were present in dividing pericycle and C_5_ cells. Cell divisions in endodermis are initiated shortly after that in C_3_ during nodule primordium development (Xiao et al., 2014). Fig. 5B shows a primordium at stage III, in which cell divisions have occurred in C_3_, but not in endodermis yet. However, *MtHDT2* transcripts were detected in nuclei of endodermal cells, indicating that *MtHDT2* starts to express in cells prior to division.

**Figure 5.**
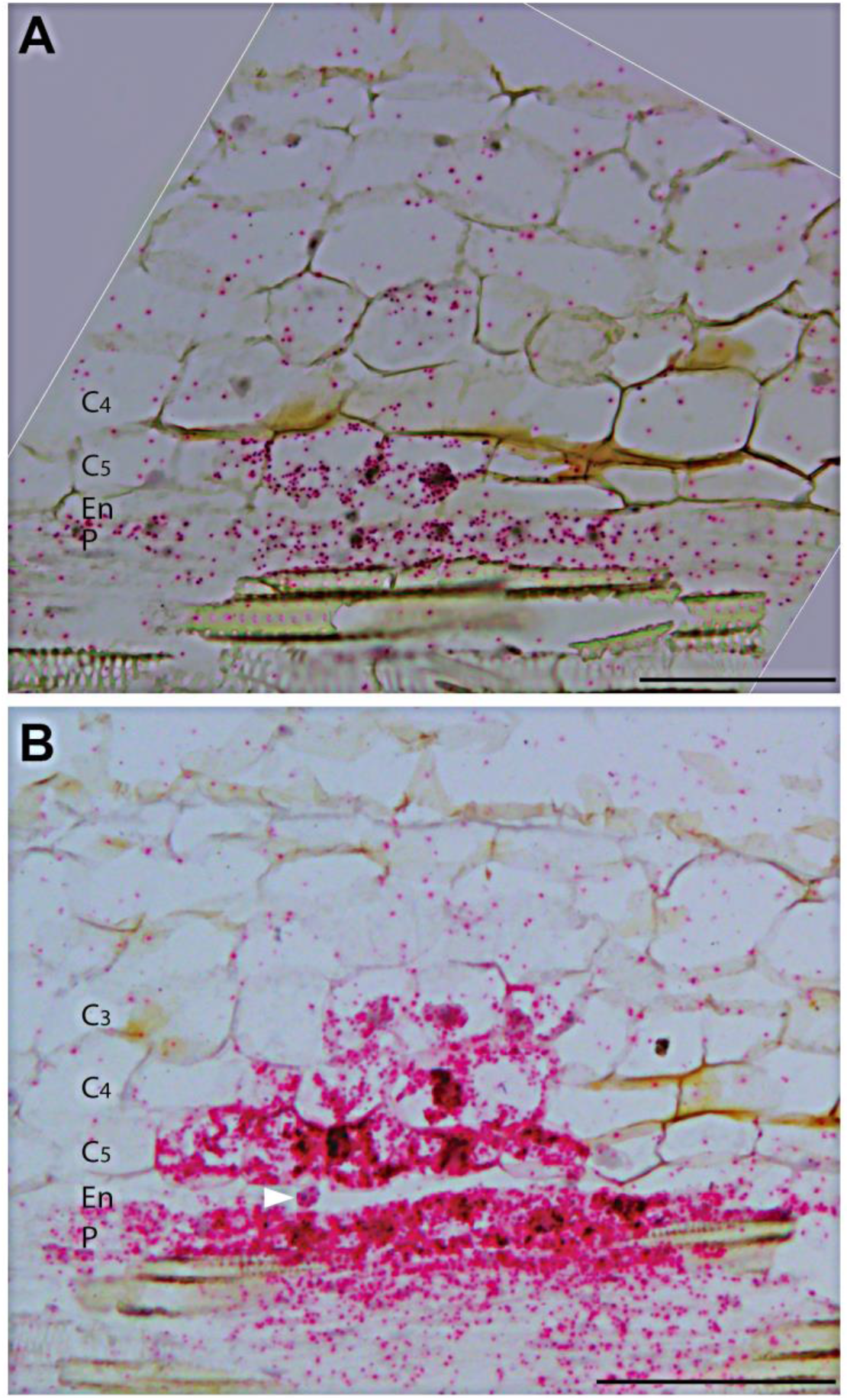
*MtHDT2* is expressed in nodule primordia. *In situ* hybridisation pattern of *MtHDT2* mRNA in nodule primordia at stage I (A) and stage III (B). Longitudinal sections of wild-type nodule primordia are shown. Red dots are hybridisation signals. Divided and dividing primordium cells are distinguished by their small size. Arrowhead in B indicates a nucleus from an endodermal cell that has not divided. P, Pericycle; En, Endodermis; C_5/4/3_, the fifth/fourth/third cortical cell layer. Scale bar=100μm.

As the expression level of *MtHDT1*, *3* is rather low, we were not able to study their expression in primordia with *in situ* hybridisation. Therefore, the expression patterns of *MtHDT1* and *MtHDT3* in nodule primordia was studied by using promoter-GUS constructs. The *pMtHDT1::GUS* construct was generated by fusing the *MtHDT1* putative promoter with *GUS* coding region and the *pMtHDT3::GUS* construct was as aforementioned. These two constructs were introduced into Medicago by hairy root transformation and transgenic roots were inoculated with rhizobia. We first analysed their expression pattern in nodules. The GUS expression patterns were consistent with RNA *in situ* hybridisation (Figs. 2, C and D; Supplemental Figs. S5, A and B), indicating that the putative promoters are sufficient to create the correct gene expression pattern. In nodule primordia, both *MtHDT1* and *MtHDT3* promoters showed a similar expression pattern as *MtHDT2* (Supplemental Figs. S5, C and D). The expression of *MtHDTs* in young primordia indicates that they have a role in nodule primordium initiation and development.

### Knock-down of*MtHDTs* Alters Gene Expression in Nodules

HDT proteins are known to regulate chromatin status by which they contribute to the regulation of transcription of genes (Kouzarides, 2007). To investigate which genes are regulated by MtHDTs, RNA-seq analyses were conducted. We isolated RNA from nodules, as it was not well possible to collect sufficient primordium material and especially because the majority of the *MtHDTs RNAi* primordia were blocked in development, this might have caused secondary effects. We collected apical part of nodules including meristem and infection zone as *MtHDTs* are preferentially expressed there. To dissect them from the fixation zone, transgenic control and *MtHDTs RNAi* roots were inoculated with rhizobia expressing *nifH::GFP*. The *nifH* gene is switched on at the transition from infection to fixation zone (Gavrin et al., 2014), where *MtHDTs* are also switched off. We will name the part, containing meristem and infection zone, nodule apex.

Transcriptomes of control and *MtHDTs RNAi* nodule apices were analysed and we detected the transcripts of ~20,000 genes in each sample (Supplemental Dataset S1). The reduced expression level of *MtHDTs* and *MtPLT3,4* in *MtHDTs RNAi* nodule apices is consistent with qRT-PCR data (FigS.3, A and C; Supplemental Dataset S1), indicating that RNA-seq data are reliable. To identify differentially expressed genes (DEGs), we performed relatively stringent statistics and filtering (fold change>4 and FDR p-value<0.05). In total 49 DEGs were identified between control and *MtHDTs RNAi* nodule apices (Supplemental Dataset S1).

To investigate whether HDTs control nodule development by regulating the same genes as in Arabidopsis roots, we first checked the expression of *GA2ox* genes as they are targets of HDTs in Arabidopsis roots (Li et al., 2017). However, *MtGA2ox* genes, were not among the 49 DEGs (Supplemental Dataset S1), suggesting that HDTs regulate nodule and root development in a different way. To further test this, we compared the DEGs that are identified in Medicago nodule apices (n=49) with those of Arabidopsis root tips (n=217) (Li et al., 2017). Gene orthology of the two species is well studied (van Velzen et al., 2018). 63% (31 out of 49) of the Medicago DEGs have (an) orthologous gene(s) in Arabidopsis, but only the 2 *HDT* genes (*MtHDT1/2*, *AtHDT1/2*) were down-regulated in both RNAi experiments (Supplemental Dataset S2). This demonstrated that none of the DEGs, that is the result of down-regulation of HDTs, was in common in Arabidopsis roots and Medicago nodules. We concluded that HDTs regulate nodule and root development in a different way.

To obtain insight in the biological functions of the identified 49 DEGs from nodule apices, we performed Gene Ontology (GO) analysis. This showed that genes encoding proteins with terpene synthase, methyltransferase or oxidoreductase activities were enriched among the DEGs (Supplemental Fig. S6).

### MtHDTs Possibly Control Nodule Development by Regulating *MtHMGR1* Expression

Two DEGs encode 3-hydroxy-3-methylglutaryl-coenzyme A reductases (*MtHMGR1* and *MtHMGR4*). These two genes were down-regulated 8.7 (*MtHMGR1*) and 7.7 (*MtHMGR4*) fold in *MtHDTs RNAi* nodule apices, respectively (Supplemental Dataset S1). Previously, it has been shown that knock-down of *MtHMGR1* blocks nodule formation (Kevei et al., 2007). The function of MtHMGR1 in mature nodules has not been studied, but it has been shown to be an interactor of MtDMI2 (Kevei et al., 2007). Knock-down of *MtDMI2* in nodules affects the intracellular colonization of rhizobia (Limpens et al., 2005), similar to that in *MtHDTs RNAi* nodules. Therefore we focused on *MtHMGR1*.

To determine in which tissue *MtHMGR1* is expressed and whether knock-down of *MtHDTs* affects its expression pattern, we performed RNA *in situ* hybridisation on longitudinal sections of nodules harvested at 21dpi. In control nodules, *MtHMGR1* was expressed in nodule meristem and the infection zone, in the latter its expression only occurred in the infected cells (Fig. 6A). In *MtHDTs RNAi* nodules, *MtHMGR1* had the same expression pattern (Fig. 6B), albeit at a markedly lower level (Supplemental Dataset S1).

**Figure 6.**
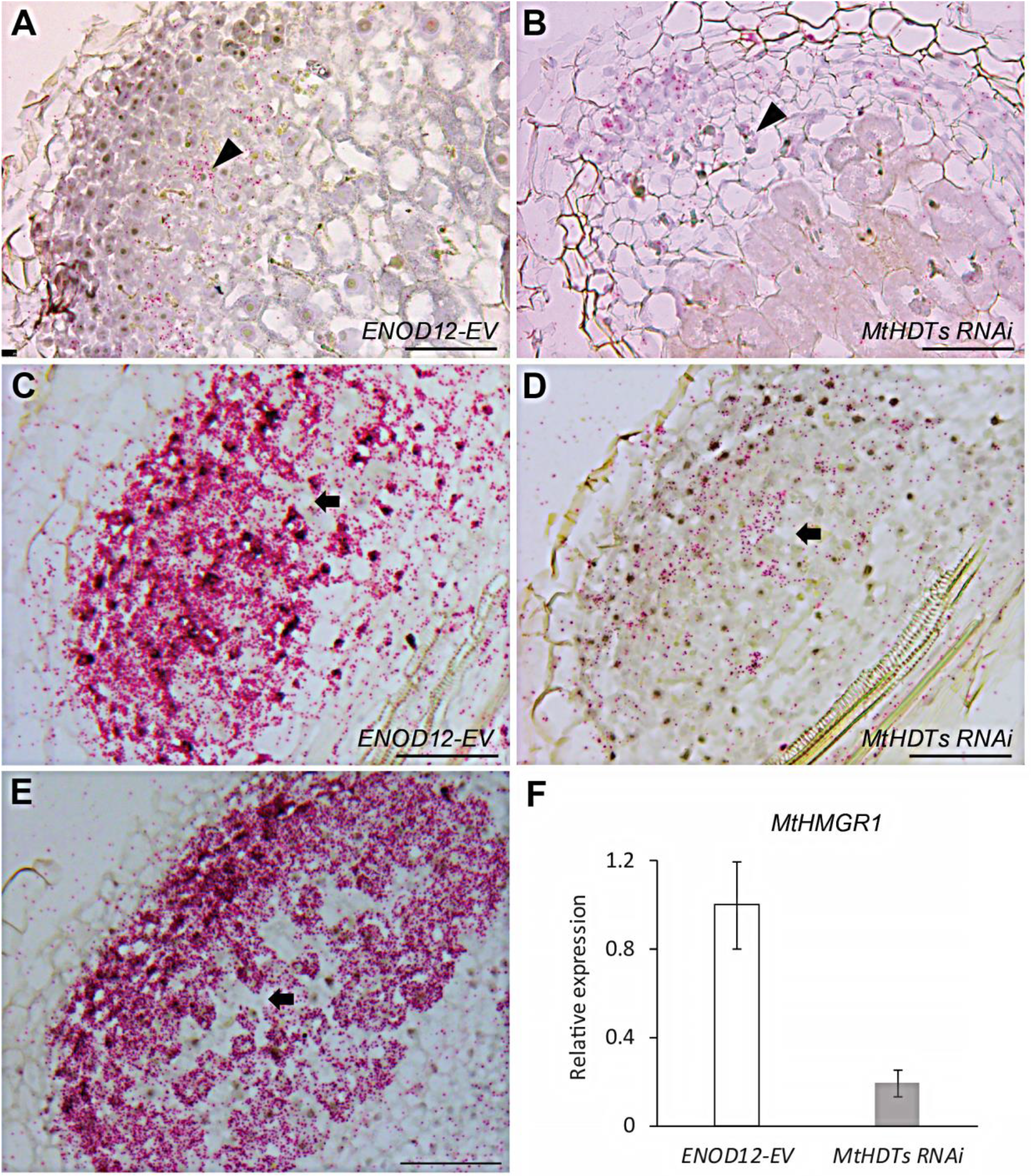
MtHDTs regulate the expression of *MtHMGR1*. A and B, *In situ* hybridisation pattern of *MtHMGR1* mRNA in *ENOD12-EV* (A) and *MtHDTs RNAi* (B) nodules. Arrowheads indicate infected cells in the infection zone. C and D, *In situ* hybridisation pattern of *MtHMGR1* mRNA in *ENOD12-EV* (C) and *MtHDTs RNAi* (D) nodule primordia. Arrows indicate non-infected cells. E, *In situ* hybridisation pattern of *MtHDT2* mRNA in wild-type nodule primordium. The arrow indicates a non-infected cell. F, RT-qPCR analysis of *MtHMGR1* expression in *ENOD12-EV* control and *MtHDTs RNAi* nodule primordia. Data shown is mean±SEM determined from three independent experiments. Nodules and nodule primordia were harvested at 21dpi and 5dpi, respectively. In A to E longitudinal sections of nodules (A and B) or nodule primordia (C to E) were shown. Red dots are hybridisation signals. Scale bar=100μm.

It has been shown that knock-down of *MtHMGR1* blocks nodule primordium development, similar to the phenotype of the inoculated *MtHDTs RNAi* roots. We then asked whether the expression pattern and level of *MtHMGR1* in nodule primordia was affected by knocking-down of *MtHDTs*. To answer this, RNA *in situ* hybridisation with *MtHMGR1* probe set was performed on longitudinal sections of 5dpi nodule primordia. In control nodule primordia (stage V), *MtHMGR1* transcripts were very abundant in (future) meristem and infected cells (Fig. 6C). Expression pattern of *MtHMGR1* in *MtHDTs RNAi* nodule primordia (stage V) resembled that of the control (Fig. 6D) albeit at a reduced level. qRT-PCR confirmed this reduced expression level (Fig. 6F). This is in line with the observation in mature nodules where knock-down of *MtHDTs* does not affect *MtHMGR1* expression pattern, but only reduced its expression level (Figs.6, A and B; Supplemental Dataset S1).

In nodules the expression pattern of *MtHMGR1* coincides with that of the *MtHDTs* (Fig. 2; Figs. 6, A and B). To test whether in nodule primordia *MtHMGR1* and *MtHDTs* were expressed in the same cells as well, we performed RNA *in situ* hybridisation with *MtHDT2* probe set on longitudinal sections of 5dpi nodule primordia. This revealed that in nodule primordia (Stage V) *MtHDT2* was also expressed in the future nodule meristem and infected cells (Fig. 6E), similar to *MtHMGR1*.

Taken together, our data showed that *MtHDTs* and *MtHMGR1* were co-expressed during nodule development. Knock-down of *MtHDTs* did not affect the expression pattern of *MtHMGR1*, but only its expression level.

## DISCUSSION

In this study, we showed that the MtHDTs play a key role in both nodule primordium formation and nodule development. Knock-down of *MtHDTs* caused a block of primordium development and in nodules it reduced meristem size and rhizobial colonization of cells. In both cases these chromatin remodelling factors positively regulate the expression of *MtHMGR1* that previously has been shown to be essential for nodule primordium formation (Kevei et al., 2007). The similar nodule primordium phenotype in *MtHDTs* and *MtHMGR1* knock-down indicates that the decreased expression of *MtHMGR1* is sufficient to explain the arrested nodule primordium development in *MtHDTs RNAi*. The mechanism by which they control nodule (primordium) development is different from that involved in root development.

We did not study the role of MtHDTs in Medicago root development. However, it seems probable that their function is similar to that of AtHDT1/ 2 in Arabidopsis roots. Firstly, this conclusion is supported by the fact that the MtHDTs and AtHDT1 and 2 have the same expression pattern in roots. Secondly, *pMtHDT2::GFP::HDT2* is sufficient to restore root development in an Arabidopsis *hdt1hdt1hdt2hdt2* background. AtHDT1, 2 regulate root meristem size by repressing *AtGA2ox2* (Li et al., 2017).

Therefore it is very probable that MtHDT2 has a similar function when expressed in Arabidopsis and it is likely that MtHDTs control Medicago root growth in a similar manner. If this is indeed the case, the mechanism by which MtHDTs regulate nodule meristem size is different, as expression of *MtGA2oxs* is not affected in *MtHDTs RNAi* nodule apices. Further none of the Arabidopsis orthologues of the Medicago nodule DEGs is affected in the Arabidopsis *HDTs RNAi* roots. In addition, the expression pattern of the *MtHDTs* in nodules and roots is not similar. In nodules the *MtHDTs* are expressed at equal levels in meristem and infection zone. The latter is equivalent to the root elongation zone. However, in roots the *MtHDTs* are expressed at the highest level in the meristem, whereas in the elongation zone their expression level is very low.

It has been shown that the nodule and root developmental programmes share transcription factors like PLETHORA and LBD16. It is possible that during the development of these two organs different genes are regulated by these transcription factors. For example during nodule development LBD16 interacts with a CCAAT box-binding protein Nuclear Factor-Y (NF-YA1), the latter is a nodule-specific transcription factor. The expression of *LBD16* is directly regulated by NODULE INCEPTION (NIN) (Schiessl et al., 2019; Soyano et al., 2019). NIN is nodule-specific transcription factor as well (Combier et al., 2006; Marsh et al., 2007), indicating that the expression of *LBD16* is also regulated differently during the development of both organs. Further, 96% of the transcriptional changes are shared with *nin* and *lbd16* loss-of-function mutants. It is probable that the genes regulated by LBD16 during the development of both organs are not completely identical. Our study shows that chromatin remodelling factors *HDTs* are involved in root and nodule development, and their targets in these two processes are also different. So although root and nodule development share several regulators, it is possible that they have different functions.

Another chromatin remodelling factor, DNA demethylase (*MtDME*) has been shown to be expressed in nodule infected cells. Knock-down of this gene does not decrease nodule number, but reduces the endoreduplication level of infected cells (Satge et al., 2016). *MtDME* is expressed at a low level in roots and its role in root development has not been studied. So whether it has a similar function in roots and nodules is unknown. During nodule development, *MtDME* first becomes active when rhizobial infection into cortical cells has already taken place. We show that *MtHDTs* are induced much earlier than *MtDME*, since the expression of *MtHDTs* is detected in nodule primordium cells prior to division (Fig. 5; Supplemental Fig. S5). Similar to this, during initiation of lateral root primordium, *AtHDT1/2* are induced in founder cells before the first cell division occurs (De Smet et al., 2008), suggesting that HDTs control the organogenesis of the two lateral organ primordia from the start.

It is well possible that more chromatin remodelling factors are shared between root and nodule development. Except *HDTs*, another 5 chromatin remodelling genes are up-regulated in Arabidopsis early lateral root primordium and their orthologs are up-regulated in Medicago roots inoculated with rhizobia (Supplemental Table S2) (Benedito et al., 2008; De Smet et al., 2008), it will be worthwhile to compare their function during root and nodule development.

Knock-down of the 3 *MtHDTs* resulted in a nodule phenotype, whereas the only available Medicago *hdt2* single mutant makes WT-like nodules, suggesting a functional redundancy of MtHDTs. Similarly, in Arabidopsis, both AtHDT1 and 2 control root development and leaf polarity (Ueno et al., 2007; Li et al., 2017). The redundancy might be due to the fact that *AtHDTs* as well as *MtHDTs* are the result of a recent gene duplication. In some monocots such duplication has not occurred (Pandey et al., 2002; Grandperret et al., 2014), and knock-down of a single *OsHDT701* (Fig. 1A) gene in rice enhances resistance to pathogens (Ding et al., 2012).

Silencing of *MtHDTs* resulted in a block of nodule primordium formation. We used the *ENOD12* promoter to silence the *MtHDTs*. During nodule primordium initiation the activation of this promoter could only be detected in pericycle and inner cortex when cell division has already occurred (Supplemental Fig. S7). This implies that most likely silencing is first effective when primordium formation has already been initiated. Therefore it is well possible that MtHDTs are essential from the start of primordium initiation. In Arabidopsis roots, silencing of *AtHDT1,2* does not affect progression through the cell cycle. However, in most nodule primordia present in *MtHDTs RNAi* roots DNA synthesis is blocked or markedly reduced indicating that cell division is (getting) blocked in these primordia (Fig. 4). This further supports that HDTs have different roles in root and nodule development.

Although the MtHDTs are important for primordium development, still a few nodules were formed on *MtHDTs RNAi* roots. Most likely in these cases expression of *MtHDTs* is not sufficiently reduced to block primordium development. In mature Medicago nodules the ~8 proximal cell layers with infected cells are derived from the primordium and not from the meristem (Xiao et al., 2014). In the few nodules formed on *MtHDTs RNAi* roots, rhizobial colonization is not affected in these infected cells derived from the primordium, but it is strongly reduced in cells of the infection zone derived from the nodule meristem. This difference in efficiency of colonization is in agreement with the idea that rhizobial infection in nodule cells is more stringently controlled than in primordium cells (Combier et al., 2006; Laporte et al., 2014).

The expression of *MtHDTs* in nodule meristem and infection zone is consistent with their function in colonization of infected cells, as well as in specifying nodule meristem properties. Considering that the *MtHDTs RNAi* nodule meristem is smaller, the reduced colonization in cells derived from this meristem might be the indirect effect of altered properties of the meristem cells. The cells of the meristem of these nodules still divide, whereas *MtHDTs RNAi* results in a block of cell division in primordia. However, the nodules are formed from primordia in which cell division is not (fully) blocked, most likely due to less reduction of the *MtHDTs* mRNA levels.

At the transition from infection to fixation zone, *MtHDT2* expression level dropped dramatically (Fig. 2A). At this transition several other sudden changes occur, including accumulation of starch in the infected cells, collapse of the vacuole of the infected cells and the induction of *nifH* genes of the rhizobia (Gavrin et al., 2014). So the sudden decrease of *MtHDT* transcripts and proteins supports the existence of a molecular switch at this transition.

Expression patterns of *MtHMGR1* and *MtHDTs* overlapped in both nodule primordia and nodules (Figs. 2 and 6), indicating that MtHDTs regulate *MtHMGR1* expression in a cell autonomous manner. *MtHMGR1* is down-regulated in *MtHDTs RNAi* primordia as well nodules and the transcriptome studies shows that all other *MtHMGR* members are down-regulated in *MtHDTs RNAi* nodule apices (Supplemental Dataset 1). They encode enzymes that catalyse the rate-limiting step in the mevalonate pathway. This pathway leads to the synthesis of sterols and isoprenoids, that give rise to several plant hormones, for example cytokinin, gibberellin and abscisic acid (Chappell et al., 1995). Whether the disturbed isoprenoid biosynthesis results in the *MtHDTs RNAi* phenotype cannot be excluded.

However, it has also been shown that *MtHMGR1* knock-down affects Nod factor signalling as it blocks rhizobium induced Ca^2+^spiking in the epidermis (Venkateshwaran et al., 2015). As Nod factor signalling is required for nodule primordium formation this function can explain the primordium phenotype in both *MtHMGR1* and *MtHDTs RNAi* (as in the latter the expression of *MtHMGR1* is reduced). Nod factor signalling also occurs in the distal part of the infection zone. Knock-down of Nod factor receptor genes as well as an essential component of the Nod factor signalling cascade *DMI2* results in reduced colonization of rhizobia in nodule cells (Limpens et al., 2005; Moling et al., 2014). This phenotype is similar to that of the *MtHDTs RNAi* nodules. So in case MtHMGR1 is required for Nod factor signalling at early stages as well as in the nodule its reduced expression can explain the *MtHDTs RNAi* nodule primordium and nodule phenotypes.

## MATERIALS AND METHODS

### Plant Growth, Transformation and Rhizobial Inoculation

*Medicago* ecotype Jemalong A17 and *ENOD11::GUS* stable line (Journet et al., 2001) were used in this study. *Agrobacterium rhizogenes* MSU440 mediated hairy root transformation was performed according to (Limpens et al., 2004). The composite plants with transgenic roots were grown either in perlite saturated with low nitrate containing Farhaeus medium (Fahraeus, 1957), or on plates with agarose-based BNM medium (Ehrhardt et al., 1992), at 21°C in a 16 h : 8 h, light : dark regime. *Sinorhizobium meliloti* 2011 or *S. meliloti* expressing *nifH:GFP* (Gavrin et al., 2014) liquid cultures were treated with 10μM luteolin for 24 hours, and then used to inoculate Medicago roots. Mature nodules were harvested at 21 days post inoculation (dpi) from roots of Medicago plants growing in perlite. Nodule primordia were harvested at 2 or 5 dpi from spot inoculated roots of Medicago plants growing on plates.

### Phylogenetic Tree Construction

Gene accession number of *HDTs* are shown in Supplemental Table S1. For phylogenetic reconstruction, protein sequences were first aligned using MUSCLE (Edgar, 2004) implemented in Geneious Prime 2019 (New Zealand) using default parameters. After manual inspection, geneious tree builder was applied to generate the phylogeny by using Neighbor-Joining methods (Saitou and Nei, 1987).

### Constructs

N-terminal fusions of MtHDTs with GFP under the control of their own promoters were constructed using MultiSite Gateway Technology (Thermo Fisher Scientific). The coding sequence (CDS) and putative promoter of each *MtHDT* were first PCR amplified by using Phusion high-fidelity DNA polymerase (Finnzymes) and nodule cDNA and genomic DNA were used as templates. The obtained PCR fragments were introduced into a pENTR-D-TOPO vector (Invitrogen). Each of the *MtHDT* promoters was cut out of the pENTR-D-TOPO vector using the NotI and AscI restriction enzymes, and then ligated with a BsaI digested pENTR4-1 vector (Invitrogen) containing GFP by using T4 DNA ligase (Thermo Fisher Scientific). The final pENTR4-1 vector with the *MtHDT* promoter and GFP, the corresponding pENTR-D-TOPO *MtHDT* CDS vector and a pENTR2-3 vector containing a CaMV35S terminator were recombined into the binary destination vector pKGW-RR-MGW thereby creating *pMtHDT::GFP::MtHDT* constructs.

To create *MtHDTs RNAi* constructs, the PCR fragments of about 400-500bp for each *MtHDT* CDS were amplified and then combined by subsequential PCR steps using primers with a complementary 15 bp overhang to generate one amplicon of all 3 *MtHDTs* fragments. The final product was introduced into a pENTR-D-TOPO vector (Invitrogen) and recombined in an inverted repeat orientation into the Gateway compatible binary vector pK7GWIWG2(II) driven by nodule specific *ENOD12* promoter (Limpens et al., 2005). The control vector [(*ENOD12*::***E**mpty **V**ector* (*ENOD12-EV*)] contained no coding DNA sequence. All primers used for cloning were listed in Supplemental Table S3.1 and S3.2.

### Gene Expression And RNA-Seq

Total RNA from transgenic nodules or nodule primordia was isolated using the plant RNA Easy Kit (Qiagen). cDNA was synthesized on 1μg of isolated RNA by reverse transcription with random hexamer primers using the iScript Select cDNA synthesis kit (Bio-Rad) according to the manufacturer’s instructions. Quantitative real-time PCR was performed in a 10 μl reaction system with SYBR Green super-mix (Bio-Rad). Ubiquitin was used as a reference gene. Primers used for quantitative real-time PCR are listed in Supplemental Table S3.3.

For RNA-Seq analyses, nodule meristem and infection zone were distinguished from the fixation zone under a fluorescent stereomacroscope (Leica) and manually dissected. Three independent experiments were conducted. Total RNA was extracted as described above. RNA was sequenced at BGI Tech Solutions (Hong Kong) using Hiseq2000 instrument. Sequencing data were analysed by mapping to the Medicago genome using CLC Genomics Workbench (Denmark). Gene expression levels were determined by calculating the RPKM (Reads Per Kilobase per Million mapped reads). Differentially expressed genes (DEGs) are defined based on relatively stringent statistics and filtering (fold change>4, FDR P value<0.05) within the CLC. GO enrichment analyses was performed using agriGO v2.0 (Tian et al., 2017).

### RNA *in situ* Hybridisation

The nodules and nodule primordia were fixed with 4% paraformaldehyde mixed with 5% glutaraldehyde in 50 mM phosphate buffer (pH 7.4) and embedded in paraffin (Paraplast X-tra, McCormick Scientific). Sections of 7 μm were cut by RJ2035 microtome (Leica). RNA *in situ* hybridisation was performed using Invitrogen ViewRNA ISH Tissue 1-Plex Assay kit (Thermo Fisher Scientific) according to the manual protocol (https://www.thermofisher.com/document-connect/document-connect.html?url=https%3A%2F%2Fassets.thermofisher.com%2FTFS-Assets%2FLSG%2Fmanuals%2FMAN0018633_viewRNA_ISH_UG.pdf&title=VXNlciBHdWlkZTogVmlld1JOQSBJU0ggVGlzc3VlIEFzc2F5). RNA ISH probe sets were designed and produced by Thermo Fisher Scientific. Catalogue numbers of probe sets are the following: for *MtHDT1* is VF1-14234, for *MtHDT2* is VF1-18132, for *MtHDT3* is VF1-6000218 and for *MtHMGR1* is VF1-20373. Any probe set was omitted for a negative control. Slides were analysed with an AU5500B microscope equipped with a DFC425c camera (Leica).

### EdU Staining

The composite plants with *ENOD12-EV* or *MtHDTs RNAi* transgenic roots were grown on BNM plates and spot inoculated with *S. meliloti* 2011 as described above. After 2 days, the inoculated root segments (~0.3cm) were submerged in liquid BNM medium with extra 1g/L D-glucose and were co-incubated with 10μM EdU stock for 2 hours on a shaker. The following washing and staining procedures were conducted according to (Kotogany et al., 2010).

### Microscopy And Imaging

Root fragments and nodules were fixed as mentioned above. After that they were washed with 0.1 M phosphate buffer 3 times for 15 min each, once with water for 15 min, and dehydrated for 10 min in 10%, 30%, 50%, 70%, 90% and 100% ethanol, and sequentially embedded in plastic Technovit 7100 (Heraeus Kulzer). Sections were made of 5μm using a microtome (RJ2035, Leica), stained with 0.05% Toluidine Blue (Sigma), mounted in Euparal (Carl Roth), and analysed with a Leica AU5500B microscope equipped with a DFC425c camera (Leica). Transgenic *pMtHDT::GFP::MtHDT* nodules and root segments were sectioned into 60μm slices by vibratome (VT1000, Leica) and mounted on slides with MQ water. All confocal images were acquired using Leica SP8 confocal laser scanning microscope (Leica, Germany). GFP and EdU signal were detected with an excitation wavelength of 488 nm and DsRed was detected with an excitation wavelength of 543 nm.

## Supplemental Material

**Supplemental Figure S1.** *MtHDT3* is expressed in root tips.

**Supplemental Figure S2.** Analysis of *Mthdt* Tnt1 mutants.

**Supplemental Figure S3.** Localization of MtHDT2 resembles that of AtHDT1,2 in Arabidopsis root tips.

**Supplemental Figure S4.** Expression pattern of *ENOD11::GUS* in nodule and lateral root primordia is different.

**Supplemental Figure S5.** *MtHDT1* and *MtHDT3* are expressed in nodules and nodule primordia.

**Supplemental Figure S6.** Gene Ontology (GO) enrichment analyses of DEGs in *MtHDTs RNAi* nodule meristem and infection zone.

**Supplemental Figure S7.** Expression pattern of *ENOD12::GUS* during nodule primordium development.

**Supplemental Table S1.** Gene accessions used in the phylogenetic analysis.

**Supplemental Table S2.** The up-regulated expression of chromatin remodelling genes in Arabidopsis lateral root primordia and Medicago nodule primordia.

**Supplemental Table S3.** Primers used in this study.

**Supplemental Dataset S1.** Gene expression map in the *ENOD12-EV* and *MtHDTs RNAi* nodule meristem and infection zone.

**Supplemental Dataset S2.** *HDTs* are the only overlapped DEGs in Medicago nodules and Arabidopsis roots.

## ACKNOWLEDGEMENTS

We thank Defeng Shen from Max Plank Institute (Cologne, Germany) for the phylogenetic construction. We would like to thank Xueyuan Leng and Nathalie Veltmaat who contributed as master students to this research.

